# Jaw Kinematics and Tongue Protraction-Retraction during Chewing and Drinking in the Pig

**DOI:** 10.1101/2020.08.22.261602

**Authors:** Rachel A. Olson, Stéphane J. Montuelle, Brad A. Chadwell, Hannah Curtis, Susan H. Williams

## Abstract

Mastication and drinking are rhythmic and cyclic oral behaviors that require interactions between the tongue, jaw, and a food or liquid bolus, respectively. During mastication, the tongue transports and positions the bolus for breakdown between the teeth. During drinking, the tongue aids in ingestion and then transports the bolus to the oropharynx. The objective of this study is to compare jaw and tongue kinematics during chewing and drinking in pigs. We hypothesize there will be differences in jaw gape cycle dynamics and tongue protraction-retraction between behaviors. Mastication cycles had an extended slow-close phase, reflecting tooth-food-tooth contact, whereas drinking cycles had an extended slow-open phase, corresponding to tongue protrusion into the liquid. Drinking jaw movements were of lower magnitude for all degrees of freedom examined (jaw protraction, yaw, and pitch), and were bilaterally symmetrical with virtually no yaw. The magnitude of tongue protraction-retraction (Tx) was greater during mastication than drinking, but there were minimal differences in the timing of maximum and minimum tongue Tx relative to the jaw gape cycle between behaviors. However, during drinking, the tongue tip is often located outside the oral cavity for the entire cycle, leading to differences in behaviors in the timing of anterior marker maximum tongue Tx. This demonstrates that there is variation in tongue-jaw coordination between behaviors. These results show that jaw and tongue movements vary significantly between mastication and drinking, which hint at differences in the central control of these behaviors.

**Summary statement:** Differences in the magnitude and timing of tongue and jaw movements and the anteroposterior positioning of the tongue during chewing and drinking demonstrate key differences in coordination of these behaviors.

## INTRODUCTION

Feeding and drinking are essential oral behaviors that provide organisms with the necessary nutrients, energy, and hydration for survival. In most mammals, mastication, or chewing, is an important component of feeding because it creates a safely swallowable bolus. Mastication involves interactions between occlusal surfaces of opposing upper and lower postcanine teeth and the food. In contrast, in adult mammals, the primary methods of active liquid ingestion – lapping, licking and sucking – are tongue- or lip-based behaviors involving no intentional interactions between the bolus and the teeth. Lapping is commonly used by mammals with incomplete cheeks whereas sucking is used by mammals with complete cheeks. During lapping, the tongue protrudes into the liquid, but the lips are not submerged (Crompton and Musinsky 2011; Reis et al., 2010; Thexton and Crompton 1989; Thexton and McGarrick 1988). When the tongue contacts a solid surface with lapping-like movements, the liquid is ingested by licking (Weijnen 1998). During sucking, the lips are completely submerged into the liquid and liquid transport is achieved through changes in intraoral pressure (Thexton et al., 1998).

Despite these fundamental differences, mastication and drinking are both accomplished by coordinated and rhythmic movements of the tongue and jaw controlled by the central and peripheral nervous systems. A central pattern generator (CPG) in the brainstem drives masticatory rhythm (Dellow and Lund 1971; Nozaki et al., 1986). The output of the masticatory CPG is modulated by feedback from the periodontal ligaments, jaw and orofacial muscle spindles, and tongue mechanoreceptors in order to correctly position food for processing and adjust force output (Lund and Kolta 2005; Lund and Kolta 2006; Takahashi et al., 2007; Trulsson 2007; Trulsson and Johansson 2002).

Although extensive modulation of the CPG adjusts movements as the food is chewed (e.g., Davis 2014; Dotsch and Dantuma 1989; Iriarte-Diaz et al., 2011; Thexton and Crompton 1989; Weijs and De Jongh 1977), gape cycles during mastication are highly rhythmic (Ross et al., 2007a,b, 2010, 2017). Similar CPGs regulating rhythmicity have been observed for licking, lapping, and sucking (Barlow 2009; Boughter et al., 2012; Nakamura et al., 1999; Travers et al., 1997), but with contributions from different cortical areas than for mastication (Iriki et al., 1988). While less studied, there is evidence to suggest that modulation of the CPG involved in drinking also occurs. For example, licking frequency in rats is influenced by experimental and environmental conditions (Weijnen 1998).

Whereas there is a general understanding of the changes in CNS connections between cortical and brainstem areas underlying the maturation from drinking in infants (i.e., suckling) to chewing (Iriki et al., 1988) as well as the kinematic changes across this shift (German et al., 1992, 2006; German and Crompton 1996, 2000; Westneat and Hall 1992), comparatively less is known about the differences and similarities between mastication and non-suckling drinking kinematics and motor control. Studies on the cat (Hiiemae et al., 1978; Thexton and McGarrick 1988, 1989) and the opossum (Crompton 1989) have compared jaw and tongue movements during mastication and lapping but only one study, on pigs, has compared mastication and sucking in behaviorally mature animals (Liu et al., 2009). This study, however, focuses specifically on tongue internal deformations rather than positional changes relative to the oral cavity.

These previous comparisons demonstrate that during mastication, the tongue positions the bolus along the toothrow for processing, usually unilaterally. When the jaw begins opening, the tongue protrudes to collect the food particles before retracting to reposition the bolus on the occlusal surface at the beginning of closing (Crompton 1989; Hiiemae et al., 1978). When lapping, the tongue also protrudes during early opening and then retracts later during opening, trapping the aliquot between the tongue and hard palate prior to the next cycle (Crompton 1989; Crompton and Musinsky 2011; Gart et al., 2015; Hiiemae et al., 1978; Reis et al., 2010; Thexton and McGarrick 1988; Thexton and Crompton 1989). During drinking in pigs, the tongue extends into the liquid with the snout immersed, suggesting that the tongue may assist during sucking to bring the liquid into the oral cavity (German and Crompton 2000; Thexton et al., 1998). Nevertheless, tongue movements serve distinct functions during these two behaviors – bolus placement and positioning within the oral cavity during mastication and bolus transport into and through the oral cavity to the oropharynx during drinking. This suggests that there may be behavior-dependent coordination patterns between the tongue and the jaw, particularly when viewed in the context of differences in jaw movements and overall gape cycle dynamics.

The goal of the present study is to compare jaw and tongue kinematics during mastication and sucking in the pig (*Sus scrofa,* Linnaeus 1758) using XROMM with additional soft tissue markers in the tongue. First, we will determine whether both behaviors use the same degrees of freedom during their respective gape cycles. Previous studies have demonstrated that two rotations, jaw pitch and jaw yaw, and anteroposterior translation (i.e., jaw protraction-retraction) are used during mastication (Brainerd et al., 2010; Menegaz et al., 2015; Montuelle et al., 2020a). Whereas jaw pitch reflects jaw opening and closing, jaw yaw reflects rotation about a vertical axis contributing to the characteristic “sidedness” of mastication. We hypothesize that both behaviors will utilize similar magnitudes of jaw pitch and anteroposterior translation, but jaw yaw will be absent during sucking because no sided interaction between the teeth and the aliquot is expected.

Second, we compare the temporal dynamics of gape cycles during both behaviors. We hypothesize that masticatory cycles will be longer and more variable than drinking cycles, reflecting the changing properties of the bolus throughout a chewing sequence. This variability is expected to extend to intracycle phases (e.g., fast closing, slow closing). Additionally, we hypothesize that the jaw opening phases of drinking cycles will be longer than those of masticatory cycles due to pronounced extraoral excursions of the anterior tongue during jaw opening.

Finally, we compare protraction and retraction movements of the tongue during chewing and drinking and relate these movements to the temporal dynamics of the gape cycle. We hypothesize that drinking involves higher magnitudes of tongue protraction-retraction than chewing in order to ingest and transport liquid to the oropharynx. However, because injury to the tongue can occur if jaw and tongue movements are not coordinated (Montuelle et al., 2019, 2020b), we hypothesize that the timing of protraction and retraction relative to the gape cycle is similar between the two behaviors and has low variability.

By comparing jaw and tongue movements during mastication and drinking, this study facilitates a better understanding of the dynamic control of oral behavior variation driven by interactions between central (e.g., CPGs, premotor cortex, sensorimotor cortex) and peripheral (e.g., orofacial mechanoreceptors) components of the nervous system. Because mammals exhibit two types of rhythmic drinking behaviors throughout their lifespan (i.e., suckling and either lapping, licking, or sucking), any similarities in the kinematics of mastication and drinking in weaned animals may indicate more overlap in some aspects of the central control of these behaviors or similarities in their modulation, despite differences in bolus properties or position.

## MATERIALS AND METHODS

### Study Design, Surgery, CT Scans, and Data Collection

Jaw movements in two 3-month-old female Hampshire-cross pigs (ID #s 20 and 21) were quantified using marker-based X-ray Reconstruction of Moving Morphology (XROMM) (Brainerd et al., 2010). In each animal, 5 to 7 radiopaque tantalum markers (Bal-Tec, Los Angeles, CA, USA, 1.6 mm diameter) were surgically implanted in the skull and jaw while animals were under isoflurane anesthesia (2-5%). An additional 17 markers were placed in the tongue, with only the anterior and posterior markers used in this study (see below). After 24-hours of recovery, biplanar fluoroscopy videos were recorded at 250fps using synchronized high-speed digital cameras (Oqus 310, Qualisys, Göteborg, Sweden) while the animals were feeding or drinking. During recording sessions, animals were offered 2cm x 2cm x 1cm cubes of apple or 475 ml of apple juice. Prior to each session, perforated metal sheets (part number 9255T641, McMaster-Carr, Robinson, NJ, USA) use for distortion correction and a custom Lego® calibration cube were imaged in each fluoroscopy view to aid in undistorting and calibrating the videos, respectively, following the standard XROMM workflow (Brainerd et al., 2010; Knorlein et al., 2016; Menegaz et al., 2015). Average radiation exposure settings were 100 kVp and 4.3 mA.

After marker implantation, the animals were CT scanned at The Ohio State University College of Veterinary Medicine (Columbus, OH, USA) on a GE Lightspeed Ultra CT scanner. These scans were used to create the bone models necessary to produce the XROMM animations. Once data collection was complete, a post-mortem CT scan was performed at Holzer Clinic (Athens, OH, USA) on a Philips Brilliance 64 scanner for the precision study. Meshes of bones from the CT scans were created in VGSTUDIO MAX 3.3 (Volume Graphics GmbH). All procedures were approved by the Ohio University Institutional Animal Care and Use Committee (protocol #12-U-009).

### XROMM Study and Data Analysis

XMALab (version 1.5.4; Knorlein et al., 2016) was used to perform calibrations, undistort the individual fluoroscopy videos for each sequence, track undistorted marker coordinates in each undistorted and calibrated fluoroscopic view, calculate 3D coordinates of each marker, and reconstruct rigid body transformations, which were filtered using a low-pass Butterworth filter with a cut-off frequency of 25 Hz. In short, the perforated metal sheet was imaged to determine distortions in the field of view whereas the calibration cube was imaged in multiple positions across the field in order to determine the camera position, orientation, and spacing. As this gives orientation and scale to the field of view, marker screen coordinates can then be translated to calibrated 3D space.

A joint coordinate system (JCS) was created in Maya (Autodesk Inc., San Rafael, CA, USA) using the CT reconstruction of the skull and jaw and then used to calculate rotations and translations of the jaw relative to the skull. All axes are perpendicular to each other with the x-axis running anteroposterior in the midline, the y-axis oriented dorsoventrally, and the z-axis oriented along the mediolateral plane running through both condyles (Figure 1A). Both a translation (T) and a rotation (R) is possible about each of these axes, creating a potential for six degrees of freedom (DoF) describing rigid body kinematics: Tx, Ty, Tz, Rx, Ry, Rz.

**Figure 1.**
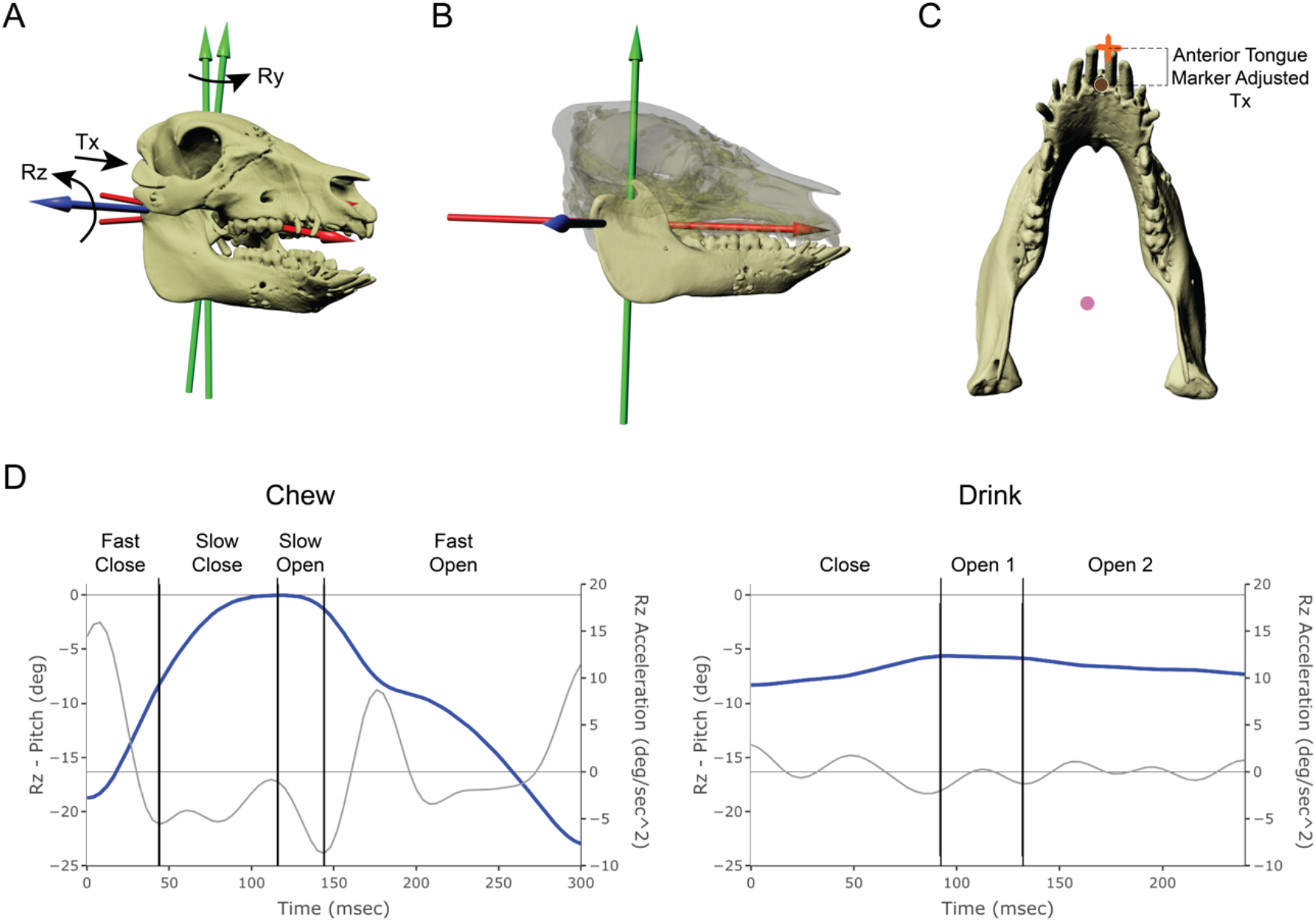
Jaw and tongue coordinate systems and tongue marker locations. (A) Orientation of the temporomandibular joint coordinate system for characterizing jaw movement, (B) orientation of the anatomical coordinate system relative to the jaw used for characterizing tongue protraction-retraction (i.e., tongue Tx), and (C) locations of the anterior (brown) and posterior (pink) tongue markers relative to the jaw at rest. Adjusted Tx values for the anterior tongue marker are corrected relative to the tip of the right lower incisor (orange X). Positive Tx indicates the anterior marker is outside the oral cavity and negative Tx indicates that the marker is inside the oral cavity. The posterior tongue marker Tx is in reference to the zero position of the jaw anatomical coordinate system shown in (B). (D) Graph of jaw pitch (Rz, blue) and acceleration (grey) during a representative chewing and drinking cycles from Individual 21 showing the differences in intracycle phases between the two behaviors. Phases for each type of cycle are based on the acceleration and directionality of Rz.

**Figure 2.**
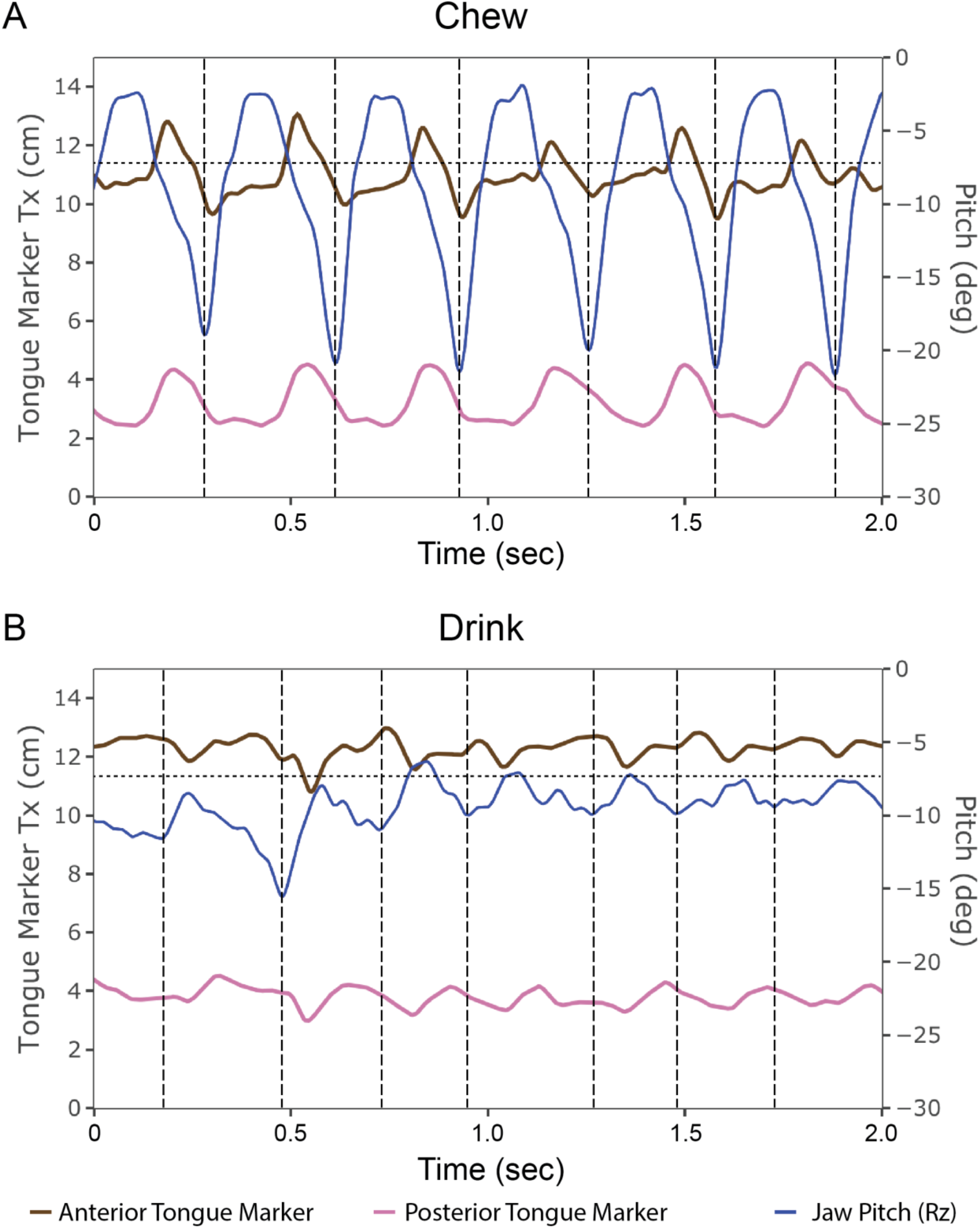
The anterior and posterior tongue markers are similar in the timing of protraction-retraction within a behavior relative to each other and relative to changes in jaw pitch. Each graph shows representative kinematic profiles of tongue marker protraction-retraction (Tx) relative to jaw pitch during chewing (A) and drinking (B) for Individual 21. The dotted horizontal line indicates the location of the incisor tip. Values above this line indicate that the anterior marker is outside the oral cavity whereas values below this line indicate that the anterior marker is within the oral cavity. Vertical dashed lines indicate minimum Rz values, or the transition between cycles.

Displacement of the tongue markers were measured relative to a jaw anatomical coordinate system (ACS) (Figure 1B). This system was a more ventrally oriented coordinate system, with the xy- and yz-planes in line with the JCS used to calculate rigid body translations and rotations, but with the xz-plane shifted dorsally so that it is positioned along the hard palate. This allows for the calculation of movements of the anterior and posterior tongue markers (Figure 1C) relative to the jaw while eliminating the influence of gape on translation in the x-dimension. Unadjusted tongue marker Tx values (anteroposterior translation: protraction-retraction) indicate displacement from the jaw ACS. Additionally, Tx of the anterior tongue marker was also adjusted so that the tip of the right central incisor defined the zero-position in the x-dimension (Figure 1C). In this context, positive Tx values indicate the anterior tongue marker being located outside the oral cavity, whereas negative Tx values indicate that it is located within the oral cavity. Note that this is only approximate as the soft tissues surrounding the oral opening (e.g., lips) are not accounted for in the rigid body motion.

After euthanasia, the frozen head of each animal was imaged within the calibrated c-arm space. Movements of the markers were then analyzed following the same XROMM workflow as above. These videos were used to calculate precision thresholds for each of the 6 DoF of rigid body motion (3 translations and 3 rotations about each of the 3 JCS axes). As no movement between the skull and the jaw is expected in the frozen specimen, any change quantified in any DoF is interpreted as digitizing error and/or error in the data collection workflow, such as suboptimal bead placement. The sequence mean of each DoF ± the precision value for each individual determines the threshold for determining jaw movements that exceed error and can be interpreted as real biological motion. Precision thresholds for each animal are provided in Supplemental Table 1.

Waves representing the DoF that exceeded the precision thresholds along with the waves representing tongue protraction-retraction were then analyzed in a custom MATLAB script (FeedCycle: Dr. Brad Chadwell, Idaho College of Osteopathic Medicine) that uses Rz (jaw pitch), the second derivative of Rz (jaw pitch acceleration), and Ry (jaw yaw) to identify key parameters of the gape cycle automatically. Individual cycles were defined from one instance of minimum Rz (i.e., maximum gape) to the following instance of minimum Rz. Within each cycle, maximum Rz (i.e., minimum gape) was used to determine the transition from jaw close to jaw open. The maximum negative value (i.e., deceleration) of the second derivative of Rz was then used to divide opening and closing into its constituent phases: fast close (FC), slow close (SC), slow open (SO), and fast open (FO). The partitioning of gape cycles into phases based on the acceleration of Rz revealed differences between mastication and drinking that impacted subsequent analysis (Figure 1D). The four standard phases were observed in chewing cycles (i.e., FC, SC, SO, and FO), whereas only three phases were detected in drinking cycles: one closing phase (C) and 2 opening phases (hereafter called O1 and O2) (Figure 1D). Because of these differences, we compared the phases between behaviors corresponding in the directionality (i.e., opening or closing) and acceleration of Rz. Thus, FC of mastication was compared to the single closing phase of sucking because of the comparable velocity of jaw closing. For opening, phases were compared based on their order of occurrence, i.e., SO and FO were compared to O1 and O2, respectively, given their presumed functionality in the context of the gape cycle.

For each cycle, total cycle duration and relative phase durations (expressed as a percentage of total gape cycle duration) were calculated. For each DoF, maximum magnitudes within each cycle and phase were calculated as the difference between the maximum and minimum values of a DoF and are reported as absolute values. Magnitudes reflect the main movements that occur within a time frame (cycle or phase) for that DoF.

In the feeding dataset used for statistical analysis, we eliminated non-chewing cycles (e.g., ingestion, stage I transport) and dropped all cycles containing a visible swallow. This resulted in 47 masticatory cycles and 40 sucking cycles for Pig 20 and 55 masticatory and 50 sucking cycles for Pig 21. All statistical analyses were performed in R version 3.6.1 (R core team 2019). On magnitude variables, we used linear mixed effects models with repeated measures, with behavior as a fixed factor and individual as the random factor using the nlme (*lme: Linear and Nonlinear Mixed Effects Models.* R package version 3.1-143) and emmeans (*emmeans: Estimated Marginal Means, aka Least-Squares Means)* packages. Additionally, in order to compare variability in cycle durations, the coefficient of variation (CV) was calculated for each cycle and phase within each sequence of mastication or sucking. Mean and variance of timing parameters were calculated with the CircStats package (*CircStats.* R package version 0.2-6). For timing parameter models, we used Bayesian circular mixed effects models with repeated measures, with behavior as a fixed factor and individual as the random factor using the bpnme function (10,000 iterations, 2,000 burn-in, 101 seed, n.lag=3) from the package bpnreg (*bpnreg: Bayesian Projected Normal Regression Models for Circular Data.* R package version 1.0.3) following the methods of Cremers and Klugkist (2018). This method produces the posterior mean, posterior standard deviation, and the 95% highest posterior density interval (HPD). HPDs (Supplemental Figure 1) are reported as the start position (as % of cycle duration) to end position, where directionality matters. Non-overlapping HPDs indicate a difference between behaviors whereas overlapping HPDs indicate a null hypothesis of no differences between behaviors cannot be rejected.

## RESULTS

### Jaw Movements and Cycle Dynamics

Jaw movements during rhythmic mastication exceed precision thresholds for only three of the six potential DoF: rotation about the z-axis (Rz: jaw pitch) and y-axis (Ry: jaw yaw), as well as translation along the x-axis (Tx: protraction-retraction) (Supplemental Figure 2). Ty and Tz occasionally exceed precision thresholds but are of much smaller magnitude than Tx and does not show a rhythmic pattern relative to the gape cycle. Instead, this most likely indicates noise above our precision threshold rather than true movement. In contrast, jaw movements during drinking cycles only exceed precision thresholds for Rz and Tx (Supplemental Figure 2). This reveals that, as hypothesized, jaw yaw (Ry) does not exceed precision values, and therefore, is not a significant movement during drinking.

Compared to sucking, the magnitudes of Rz, Ry, and Tx were significantly greater during mastication for whole cycles and each intra-cycle phase (Table 1). During both behaviors, the jaw reaches maximum Rz (i.e., minimum gape) approximately 40% into the cycle (Figure 3A). Jaw yaw (Ry) reaches a maximum just after minimum gape during mastication, at which point it resets for the next cycle by switching yaw direction (Figure 3B). In contrast, Ry lacked a discernible peak during sucking indicating that it is a bilaterally symmetrical behavior, unlike mastication. For both behaviors, jaw retraction (i.e., decreasing Tx) occurs during jaw closing whereas protraction (i.e., increasing Tx) occurs during opening (Figure 3C).

**Figure 3.**
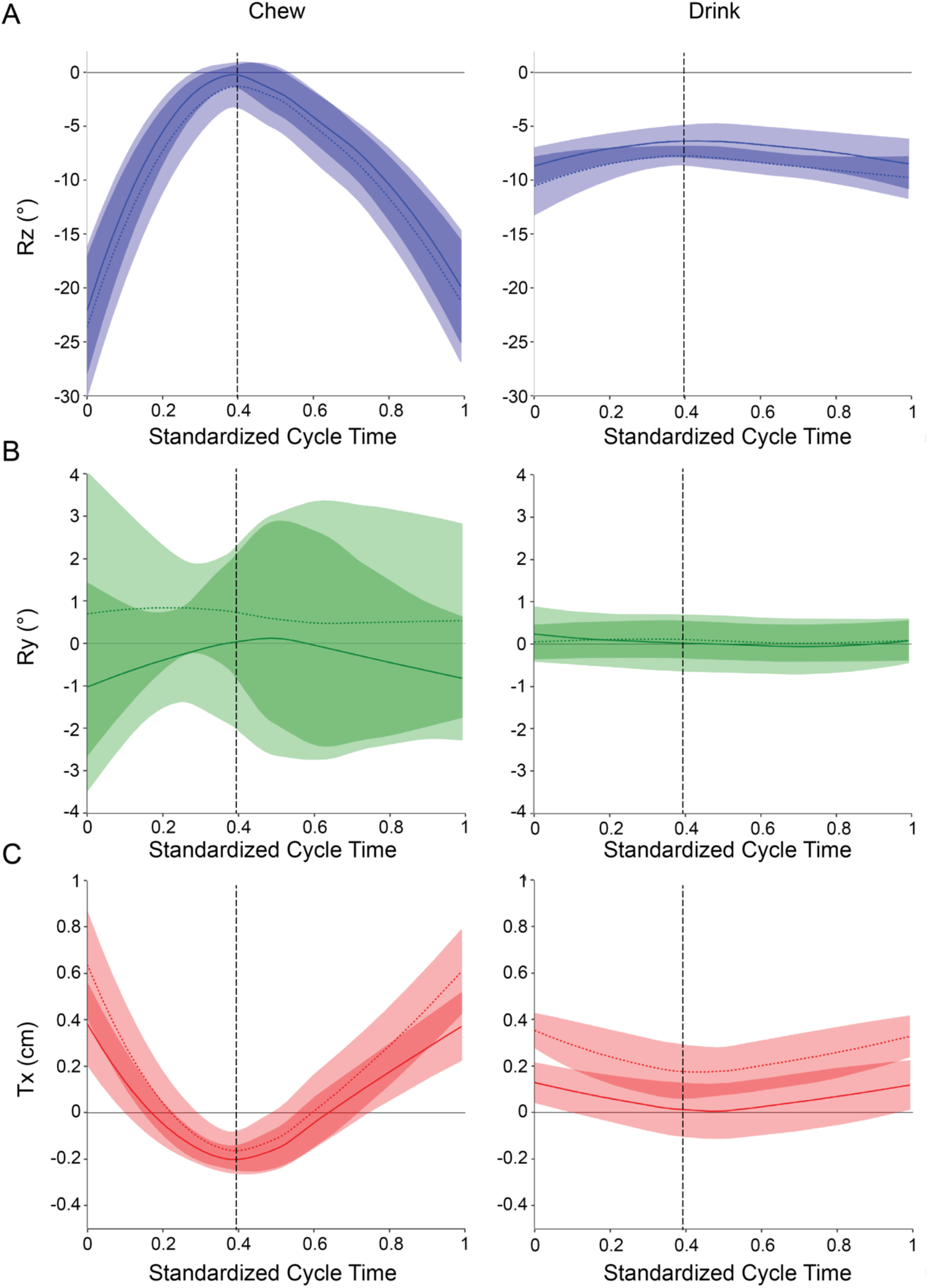
During drinking, the jaw does not reach closure and there is no appreciable yaw. Plots shows the mean and 95% confidence intervals of Rz, Ry and Tx and their 95% confidence intervals over standardized cycle times for chewing and drinking. Individual 20 is represented by solid lines and Individual 21 is represented by dashed lines. The average time of minimum gape (i.e., maximum Rz) across all cycles is indicated by the vertical dashed line.

**Table 1.** Cycle and phase level data for jaw movement and temporal dynamics during chewing and drinking cycles and corresponding model results.

|  | **Chew** | **Drink** | **Model** |
| --- | --- | --- | --- |
| **Magnitudes (Mean ± SD)** |  |  |  |
| Total Cycle |  |  |  |
| Rz (deg) | 20.8 ± 2.4 | 3.17 ± 1.2 | SE = 0.276, T<sub>2,192</sub> = 64.2, p < 0.0001 |
| Ry (deg) | 3.66 ± 0.79 | 0.708 ± 0.26 | SE = 0.0859, T<sub>2,192</sub> = 34.3, p < 0.0001 |
| Tx (mm) | 6.98 ± 1.3 | 2.08 ± 0.41 | SE = 0.117, T<sub>2,192</sub> = 42.0, p < 0.0001 |
| FC/Cl |  |  |  |
| Rz (deg) | 13.7 ± 3.3 | 2.90 ± 1.2 | SE = 0.365, T<sub>2,192</sub> = 29.6, p < 0.0001 |
| Ry (deg) | 1.32 ± 0.52 | 0.625 ± 0.28 | SE = 0.0609, T<sub>2,192</sub> = 11.4, p < 0.0001 |
| Tx (mm) | 4.29 ± 1.6 | 1.87 ± 0.59 | SE = 0.159, T<sub>2,192</sub> = 15.3, p < 0.0001 |
| SC |  |  |  |
| Rz (deg) | 6.32 ± 2.7 | — | — |
| Ry (deg) | 1.15 ± 0.79 | — | — |
| Tx (mm) | 2.25 ± 0.97 | — | — |
| SO / O1 |  |  |  |
| Rz (deg) | 6.12 ± 5.1 | 1.43 ± 0.67 | SE = 0.540, T<sub>2,192</sub> = 8.68, p < 0.0001 |
| Ry (deg) | 2.20 ± 0.97 | 0.431 ± 0.24 | SE = 0.105, T<sub>2,192</sub> = 16.9, p < 0.0001 |
| Tx (mm) | 2.68 ± 2.0 | 0.976 ± 0.61 | SE = 0.223, T<sub>2,192</sub> = 7.65, p < 0.0001 |
| FO / O2 |  |  |  |
| Rz (deg) | 13.7 ± 5.3 | 1.29 ± 0.89 | SE = 0.8564, T<sub>2,192</sub> = 22.1, p < 0.0001 |
| Ry (deg) | 1.89 ± 1.2 | 0.345 ± 0.21 | SE = 0.126, T<sub>2,192</sub> = 12.2, p < 0.0001 |
| Tx (mm) | 3.95 ± 2.3 | 0.963 ± 0.54 | SE = 0.235, T<sub>2,192</sub> = 12.8, p < 0.0001 |
| **Absolute Durations**<br>**(msecs, Mean ± SD (CV))** |  |  |  |
| Total Cycle | 323 ± 31 (9.50) | 281 ± 82 (29.1) | SE = 8.04, T<sub>2,192</sub> = 5.17, p < 0.0001 |
| FC / C | 57.7 ± 18 (30.9) | 120 ± 51 (42.3) | SE = 5.25, T<sub>2,192</sub> = -12.0, p < 0.0001 |
| SC | 78.3 ± 25 (31.8) | — | — |
| SO / O1 | 90.6 ± 57 (63.0) | 97.5 ± 52 (53.1) | SE = 7.73, T<sub>2,192</sub> = -0.930, p = 0.354 |
| FO / O2 | 97.1 ± 53 (54.2) | 65.0 ± 54 (83.4) | SE = 7.74, T<sub>2,192</sub> = 4.15, p < 0.0001 |
| **Relative Durations (%**<br>**Mean ± SD (CV))** |  |  |  |
| FC / C | 18.0 ± 5.8 (32.5) | 42.9 ± 13 (29.3) | SE = 1.39, T<sub>2,192</sub> = -17.9, p < 0.0001 |
| SC | 24.1 ± 7.4 (30.9) | — | — |
| SO / O1 | 27.7 ± 17 (59.5) | 34.5 ± 14 (41.2) | SE = 2.24, T<sub>2,192</sub> = -3.02, p = 0.0028 |
| FO / O2 | 30.1 ± 16 (53.2) | 23.1 ± 14 (62.1) | SE = 2.20, T<sub>2,192</sub> = 3.205, p = 0.0016 |

Masticatory cycles were significantly longer than sucking cycles (Table 1). Comparison of phases reveals that the absolute duration of C during sucking was significantly longer than the corresponding FC of mastication, and generally correspond to the total duration of FC+SC of mastication. Contrary to our hypothesis for jaw opening, SO and O1 absolute duration did not differ between the two behaviors, but FO was significantly longer than O2. Variability, as indicated by the CV (see Table 1), in average cycle duration across all sequences was lower for mastication (9.50) than for sucking (29.1) contrary to our prediction. At the phase level, however, opening phases were more variable than closing phases for both behaviors.

The relative contribution of each phase to total gape cycle duration also differed between the two behaviors (Table 1). Whereas C and O1 are proportionately longer for drinking cycles than FC and SO, respectively, FO had a higher contribution to total cycle duration for chewing cycles than O2 did for drinking cycles (Supplemental Figure 3). Higher variability in relative phase duration was also observed for opening phases of both chewing and drinking cycles relative to closing phases (Table 1).

### Tongue Protraction-Retraction

The timing of protraction and retraction of the anterior and posterior tongue markers is generally similar within a behavior relative to each other and relative to changes in jaw pitch but differences were observed between behaviors (Figure 4). During chewing, the anterior marker has minimal movement during jaw closing, then protracts at the start of jaw opening, followed by retraction as the jaw opens to maximum gape (Figure 4A). In contrast, the posterior marker during chewing is already in the process of retracting as the jaw begins to close from maximum gape. It then reaches minimum retraction near minimum gape, and subsequently changes direction to reach maximum protraction part of the way through opening, before it then begins to retract (Figure 4C). During drinking, the anterior tongue marker undergoes low amplitude movements, usually outside the oral cavity and may occasionally enter it before minimum gape (Figure 4B). Low amplitude movements are also observed for the posterior tongue marker (Figure 4D).

**Figure 4.**
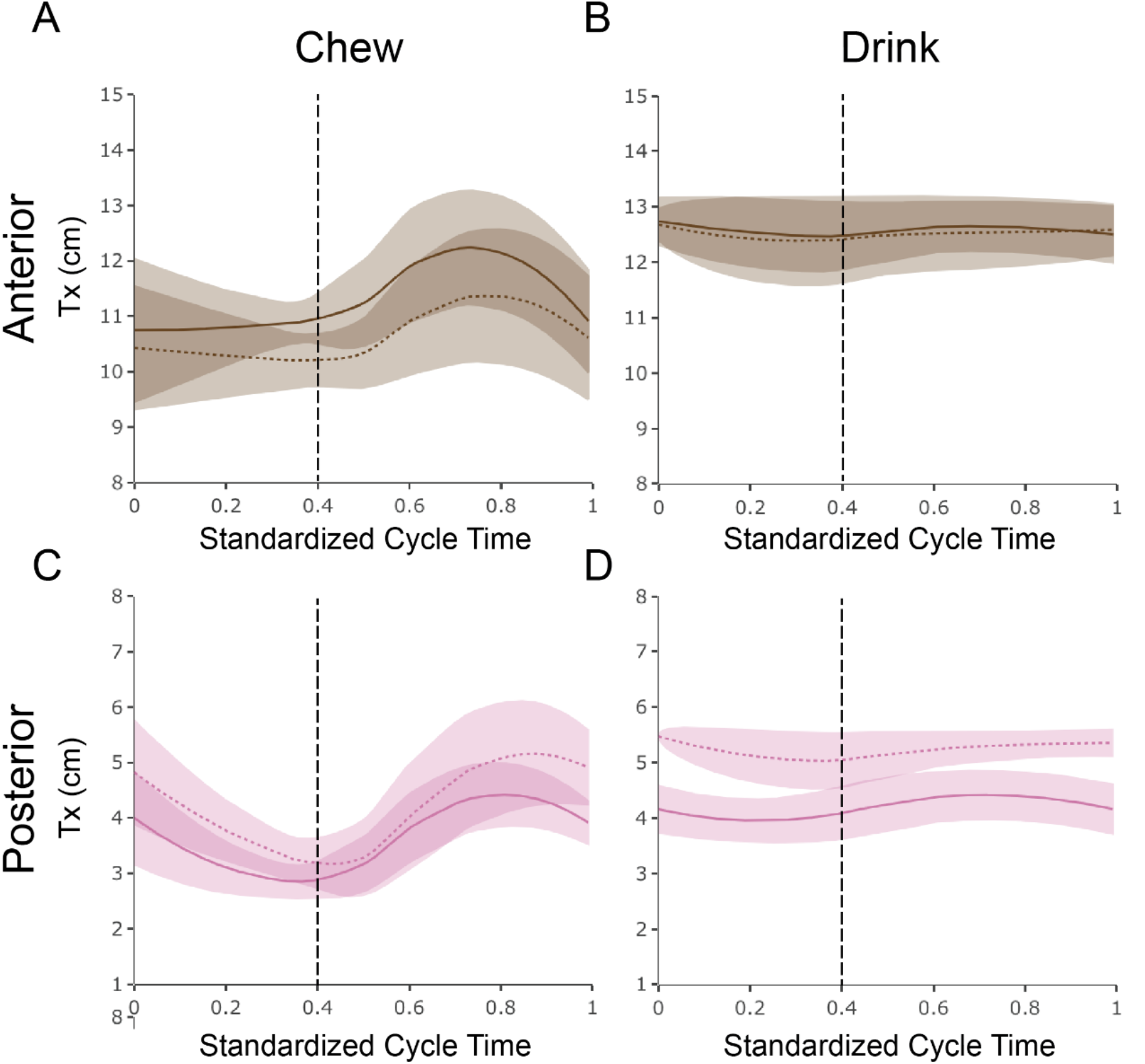
Both tongue markers undergo a greater range of protraction-retraction during chewing than during drinking. Graphs show the mean lines and their 95% confidence intervals for protraction-retraction of the anterior (A, B) and posterior (C, D) tongue markers for chewing (left), and drinking (right) plotted against standardized to cycle time. Individual 20 is indicated by solid lines and Individual 21 by dashed lines. The dashed vertical lines is the mean time of minimum gape.

Contrary to the hypothesis, Tx displacements of the anterior tongue marker are significantly larger during chewing than during drinking (Table 2; Figure 4C,D). However, the overall Tx displacement pattern of the posterior marker is more similar between behaviors than that of the anterior marker. Although Tx displacements of the anterior tongue marker are significantly smaller during drinking, the anterior tongue marker typically has significantly higher maximum and minimum Tx values during drinking compared to chewing (Table 2; Figure 4). These results indicate that the anterior part of the tongue is consistently more protracted during drinking than during chewing, and that it performs greater protraction-retraction movements during chewing. Nevertheless, maximum tongue protraction during chewing is quite variable and contains the drinking maximum and minimum protraction-retraction values within its range. The maximum protracted and retracted values of the posterior tongue marker are not significantly different between behaviors, which likely reflects regional changes in tongue deformation. However, during chewing the posterior marker retracts more than during drinking. When Tx of the anterior marker is adjusted for displacement from the lower incisor tip (Figure 5), it is clear that the anterior tongue usually protrudes outside the oral cavity during chewing then retracts into the oral cavity, whereas during drinking, it remains outside the oral cavity and only occasionally retracts back into the oral cavity. Indeed, during chewing there is usually a single excursion outside the oral cavity during jaw opening whereas during drinking, it is relatively unchanged in its position outside the oral cavity through most of the cycle (Figure 2).

**Table 2.** Anteroposterior translations of the tongue markers during chewing and drinking and corresponding model results.

|  | **Chew** | **Drink** | **Model by Behavior** |
| --- | --- | --- | --- |
| **Magnitude (mm) ± SD** |  |  |  |
| Anterior | 22.5 ± 5.9 | 6.60 ± 3.6 | SE = 0.684, $T_{2,192} = 23.2$, $p < 0.0001$ |
| Posterior | 21.2 ± 4.4 | 6.65 ± 2.0 | SE = 0.480, $T_{2,192} = 30.4$, $p < 0.0001$ |
| **Model by Marker** | SE = 0.726, $T_{2,192} = 1.73$, $p = 0.0847$ | SE = 0.425, $T_{2,192} = -0.137$, $p = 0.892$ | |
| **Maximum (mm) ± SD** |  |  |  |
| Anterior | 123 ± 6.4 | 128 ± 2.3 | SE = 0.554, $T_{2,192} = -4.64$, $p < 0.0001$ |
| Posterior | 50.5 ± 5.0 | 50.7 ± 4.5 | SE = 0.288, $T_{2,192} = -1.69$, $p = 0.0927$ |
| **Model by Marker** | SE = 0.806, $T_{2,192} = 90.5$, $p < 0.0001$ | SE = 0.450, $T_{2,192} = 172$, $p < 0.0001$ | |
| **Minimum (mm) ± SD** |  |  |  |
| Anterior | 101 ± 3.7 | 122 ± 3.5 | SE = 0.492, $T_{2,192} = -42.0$, $p < 0.0001$ |
| Posterior | 29.3 ± 1.7 | 44.1 ± 5.4 | SE = 0.372, $T_{2,192} = -39.5$, $p < 0.0001$ |
| **Model by Marker** | SE = 0.398, $T_{2,192} = 180$, $p < 0.0001$ | SE = 0.572, $T_{2,192} = 136$, $p < 0.0001$ | |

### Timing of Tongue Protraction-Retraction Relative to the Gape Cycle

The timing of maximum and minimum Tx of both tongue markers relative to the gape cycle are shown in Figure 6. During mastication, both markers reach maximum protraction during FO around 75% of the way through the gape cycle, with the anterior marker slightly preceding the posterior (Figure 6). During sucking, the anterior marker reaches its mean maximum protraction around maximum gape (i.e., at the end of O2) whereas the posterior marker reaches its mean maximum protraction earlier during O2 (Figure 6). However, the overall variance for the timing of maximum marker protraction is high, especially compared to mastication. Only the relative timing of maximum protraction of the anterior marker is statistically different between behaviors as indicated by the non-overlapping HPDs (Table 3). In contrast, HPDs for the posterior tongue marker overlap. indicating that the null hypotheses of no differences between behaviors cannot be rejected for the posterior region of the tongue. Thus, the protraction of the anterior tongue is delayed, yet more variable, during sucking compared to mastication, whereas the timing of the protraction of the posterior region of the tongue is similar during both behaviors.

**Table 3.** Results of the circular mixed effects model for the timing of maximum tongue Tx.

| | Posterior Mean ± sd<br>(start, end) | | Posterior Mean ± sd<br>(start, end) | |
| --- | --- | --- | --- | --- |
|  | Chew Cycles | Drink Cycles | Chew Cycles | Drink Cycles |
| **Anterior** | 71.1 ± 2.7<br>(65.9, 76.4) | 1.7 ± 6.7<br>(93.9, 14.2) | 8.6 ± 4.5<br>(2.5, 16.1) | 26.4 ± 6.2<br>(12.9, 40.0) |
| **Posterior** | 77.5 ± 4.5<br>(67.5, 86.8) | 86.7 ± 10.4<br>(55.1, 0.6) | 43.1 ± 3.4<br>(38.7, 46.9) | 29.2 ± 7.2<br>(13.4, 43.9) |

Values are reported as a percentage of standardized cycle time.

During mastication, the anterior tongue marker usually reaches its maximum retracted position (i.e., minimum Tx) during FC, whereas the posterior marker reaches its maximum retracted position later, usually near minimum gape (Figure 6). During drinking, both markers are usually fully retracted during closing, with relatively high levels of variance compared to chewing. This higher variance may originate from the relatively flat traces (as illustrated in Figure 4) because both locators spend a large portion of the cycle at or near their respective maximum retracted position. In spite of this variability, the anterior marker seems to be maximally retracted earlier during chewing than during drinking, whereas the reverse is true for the posterior marker (Figure 6). However, the HPD intervals for the timing of minimum Tx for both markers overlap between behaviors, indicating no statistical difference (Table 3).

## DISCUSSION

### Jaw Movements during Chewing and Drinking

As in chewing, the primary degree of freedom of jaw movements during drinking is Rz (i.e., jaw opening-closing), but the magnitude of pitch change between the two types of cycles is significantly different. Mastication requires food to be positioned and repositioned between the teeth for breakdown, necessitating a larger maximum gape during the chewing cycle. As there is no bolus between the teeth during drinking, only slight jaw opening is necessary for the tongue to protrude and retract to aid in liquid transport into and through the oral cavity. For comparison, the mean maximum Rz rotations of the jaw during chewing (−21.6°) and drinking (−9.6°) correspond to approximately 4.3 cm and 2.0 cm of gape at the incisors, respectively.

At minimum gape, the jaws almost completely close during chewing cycles, whereas during sucking cycles, the lower jaw never elevates beyond −5° (see Figure 4A) resulting in a relatively small change in jaw pitch during each cycle (3.17° ± 1.2; Table 1). Lapping in species with incomplete cheeks, such as the cat, demonstrate much larger pitch magnitudes (i.e., over 15° in the cat; Hiiemae et al., 1978) than those observed here for drinking. During lapping, the tongue is completely retracted into the oral cavity along with the water due to adhesion and inertial mechanisms, and the jaws close to pinch off the liquid column (Crompton and Musinsky 2011; Reis et al., 2010). In contrast, the low levels of jaw pitch in pigs, along with a tongue tip that often does not return to the oral cavity during drinking (Figure 5) demonstrate that in pigs, sucking is the primary mechanism of liquid transport into the oral cavity, potentially aided by small lapping-like movements of the tongue. The mechanics of sucking in relation to jaw and tongue movements are discussed further below.

**Figure 5.**
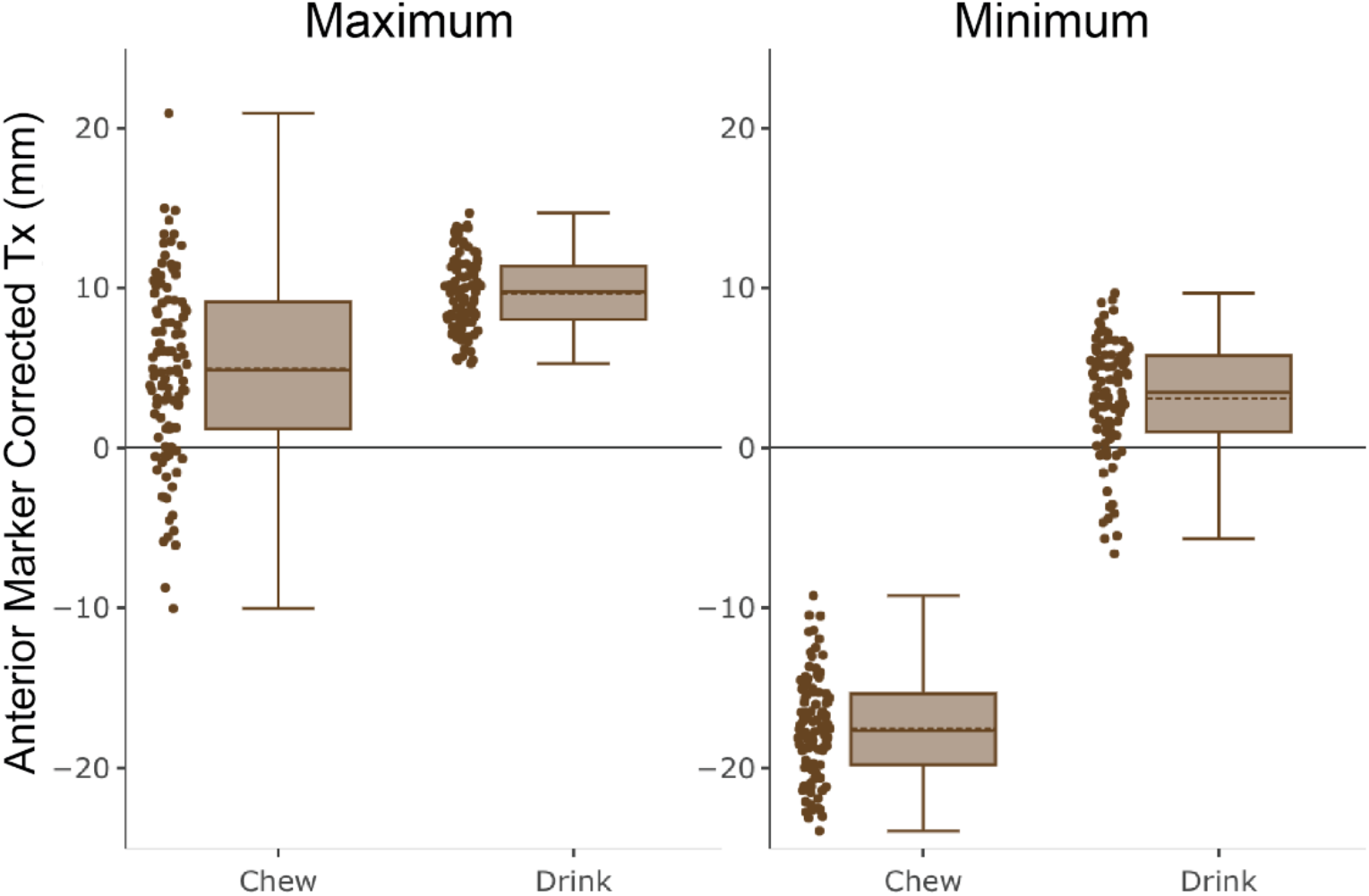
Corrected maximum Tx for the anterior tongue marker. Maximum (left) and minimum (right) Tx values for the anterior tongue marker adjusted to incisor location (see Figure 1), such that positive Tx indicates the marker is outside the oral cavity whereas negative Tx indicates it is within the oral cavity.

**Figure 6.**
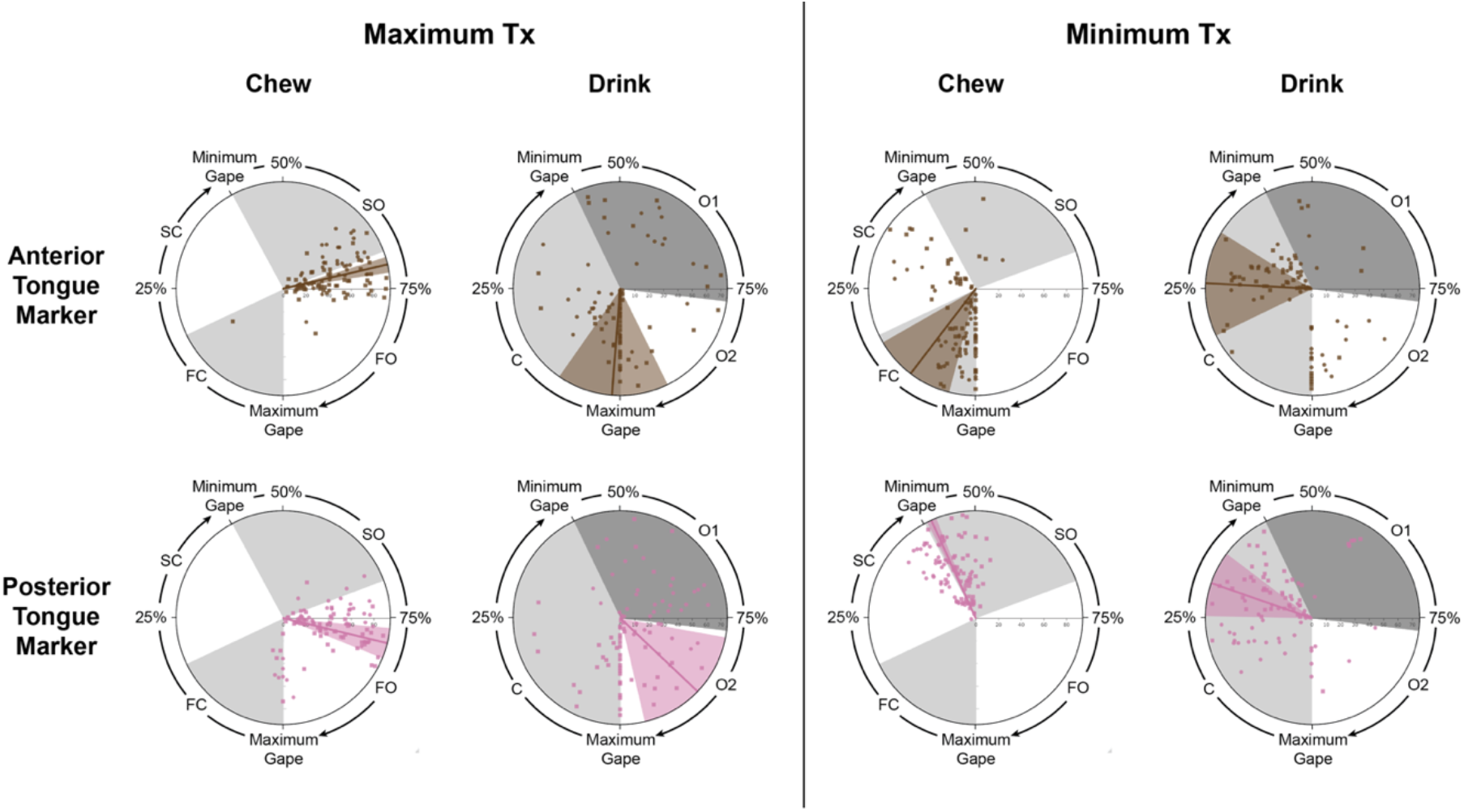
The timing of maximum Tx of the anterior tongue marker is significantly different between chewing and drinking whereas no differences are observed in the timing of the posterior tongue marker. Variance in the timing of maximum and minimum tongue Tx is typically higher during drinking than chewing. In each plot, the timing of maximum (left) or minimum (right) Tx for each tongue marker is expressed as a percent of total cycle duration and shown relative to wedges representing relative mean phase durations (alternating gray and white) during chewing and drinking. Lines indicate mean values and wedges show the corresponding variance (values reported in Supplemental Table 2). Individual 20 is indicated by circles and Individual 21 by squares. The horizontal bar indicates cycle number of the cycle from the sequence.

The other rotational degree of freedom in jaw movements during chewing cycles is yaw (Ry) to facilitate unilateral chewing. Although isognathy in pigs means that both sides occlude during a single cycle (see Herring et al., 2001), there is a clear “sidedness” to the behavior demonstrated by directionality in Ry during SC. This is supported by asymmetrical jaw muscle motor patterns in pigs despite similarities in bone strain patterns on the working- (i.e., chewing) and balancing- (i.e., non-chewing) sides (Herring 1976; Herring and Wineski 1986; Herring et al., 2001). In contrast, during drinking, changes in jaw yaw are virtually absent throughout the entire gape cycle. This confirms that drinking in pigs involves bilaterally symmetrical jaw movements, consistent with our hypothesis. Bilaterally symmetrical jaw movements also occur during infant suckling in the hamster (Lakars and Herring 1980), during food gathering and the initial cycles of nut crushing in pigs (Menegaz et al., 2015), and they can be inferred for suckling in the pig from their bilaterally symmetrical jaw muscle motor patterns (Herring 1985b).

Finally, previous work on pigs also shows that of the three available translational DoF, jaw movements during chewing cycles only use anteroposterior (Tx) translations (Brainerd et al., 2010; Menegaz et al., 2015; Montuelle et al., 2018, 2019, 2020a). We show here that this is also the case for drinking cycles. Moreover, the timing of jaw protraction and retraction is similar between the behaviors as hypothesized: jaw retraction occurs primarily during closing and protraction occurs primarily during opening as was expected. However, the magnitude of Tx is much lower during drinking because the jaw is operating over a much narrower range of pitch change. This decreases the translation necessary at the temporomandibular joint (TMJ). As Tx still exceeds the precision threshold during drinking, and there is also linear correlation between jaw pitch and protraction-retraction during both behaviors (Figure 7), this demonstrates the basic translational-rotational anatomical coupling mechanism within the TMJ that is typical of many mammals. The functional significance of this mechanism is still debated, but one likely hypothesis is that it maximizes the mechanical advantage of the masseter throughout changes in jaw pitch (Chen 1998; Hylander 1992; Smith 1985). Anteroposterior translation of the jaw may also help to align teeth and aid food breakdown during chewing. During drinking, however, tooth alignment may not be as critical because there is no tooth-food-tooth contact necessary for food breakdown.

**Figure 7.**
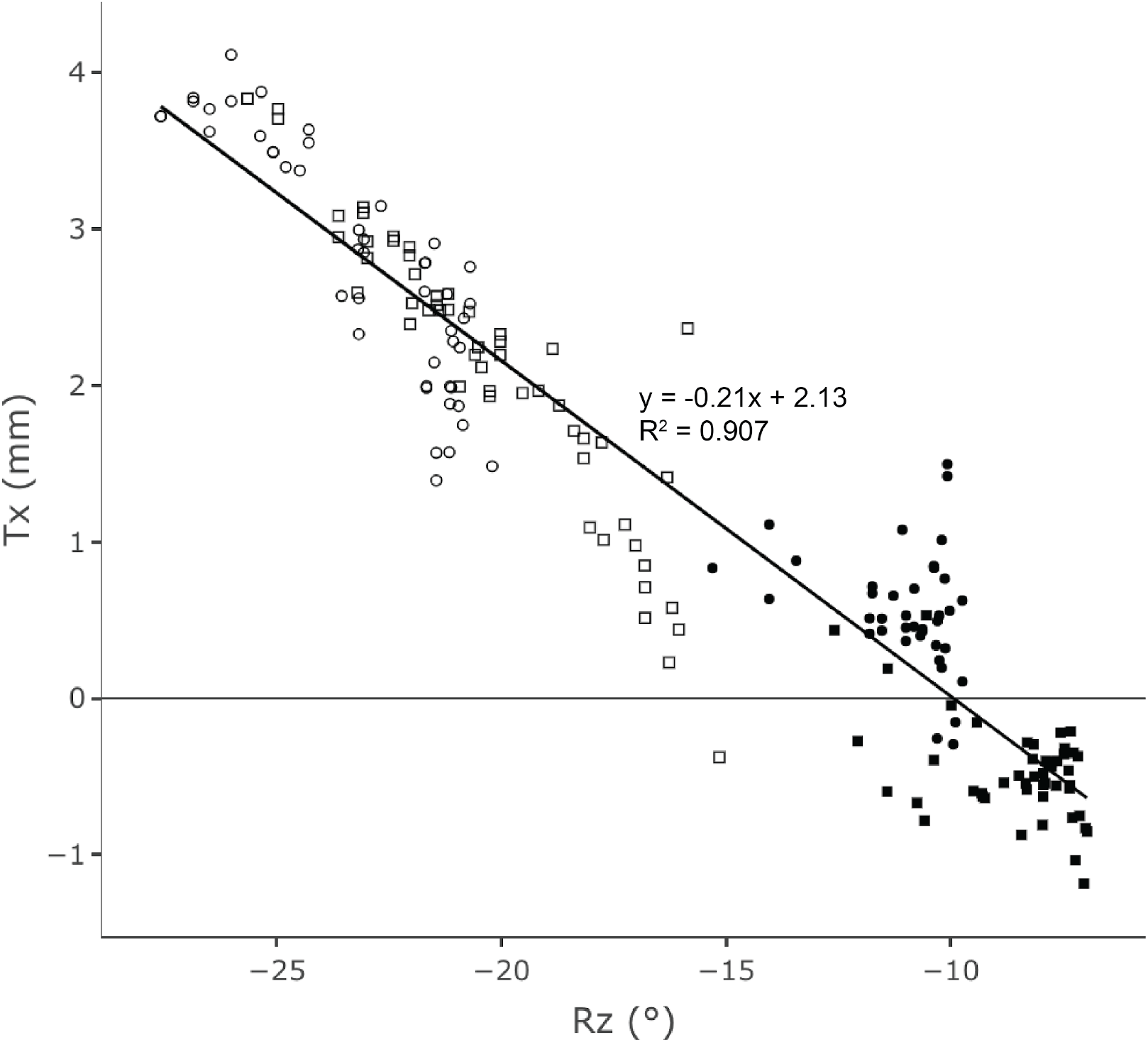
Jaw protraction-retraction (Tx) is strongly correlated with jaw pitch (Rz). This demonstrates the basic translational-rotational anatomical coupling mechanism within the TMJ that is typical of many mammals. Each datapoint represents jaw Tx and its corresponding maximum pitch (i.e., minimum Rz value) for a cycle. Chewing cycles are indicated by open symbols and drinking by solid symbols. Individual 20 is represented by circles and Individual 21 by squares. The least squares linear regression line and corresponding R^2^ is shown for combined chewing and drinking cycles.

### Cycle and Phase-Level Durations during Chewing and Drinking

We initially hypothesized that chewing cycles would be longer and more variable due to the interactions of the teeth and tongue with the food to produce a swallowable bolus. Chewing cycles were on average indeed significantly longer but, contrary to our hypothesis, less variable than drinking cycles. This may be a function of both temporal and spatial factors. Whereas chewing has an SC phase in which jaw closing slows down when the teeth contact the food, this phase was not present in drinking cycles. Rather, there is a single closing phase similar to the FC of chewing in terms of pitch velocity and acceleration. Accordingly, the absolute time spent in jaw closing was longer for chewing (see Table 1). Second, the FO phase was significantly longer during chewing cycles than its opening phase counterpart, O2, during drinking. Finally, the magnitude of jaw opening was larger during chewing and therefore, absent changes in jaw velocity through the cycle, this would extend cycle duration. However, only a weak correlation is observed in the relationship between cycle pitch magnitude and cycle duration for both behaviors (Supplemental Figure 4). Interestingly, both behaviors were similar in the relative amount of time spent during jaw closing and during jaw opening, although individual relative phase durations differed (see Table 1; Supplemental Figure 3).

The differences between chewing and drinking cycle variability are interesting in light of similar analyses from pigs and broader analysis across vertebrates. The results presented here for chewing are comparable to those reported in previous studies on chewing (e.g., Montuelle et al., 2018) and lower than those reported here for drinking. In fact, the comparatively high variability in drinking cycle duration is more consistent with that observed for lepidosaur feeding (Ross et al., 2007a, 2010). It has been hypothesized that protection of the teeth in mammals, rather than energetic savings, facilitates the low CV values observed for cycle duration across mammalian mastication (Ross et al., 2017). As drinking does not have the same constraint relating to tooth protection, there may be less constraints for a central control mechanism that maintains high rhythmicity comparable to mastication.

We hypothesized that opening phases would be longer during drinking than chewing due to extraoral excursions of the anterior tongue during these phases. Instead, total close and total open durations were similar between behaviors. There were, however, differences in the absolute and relative durations of the opening phases. The first opening phase was longer during drinking than chewing, and the second opening phase was longer for chewing (both absolute and relative). The relative duration of chewing SO decreases in pigs as food stiffness and toughness increase (Montuelle et al., 2018), such that the relationship observed in this study between chewing and drinking is likely to hold across other foods. Therefore, this long initial opening phase, in which the oral cavity increases in volume, may be functionally relevant to the creation of the pressure gradient necessary for sucking.

We also hypothesized that opening phases would be more variable during chewing than drinking because of the interactions with the food, which changes properties throughout a sequence. Variability was indeed higher for chewing than drinking cycles for both absolute and relative duration of the first opening phase, and the opposite was observed for the second opening phase. As occlusion extends into the early stages of the first opening phase (SO; Montuelle et al., 2020a), this higher variability during chewing may be attributed to the changing bolus properties.

### Tongue Protraction-Retraction

Contrary to our hypothesis, the magnitude of anterior tongue protraction and retraction is higher during chewing compared to drinking. During most chewing cycles, the anterior tongue marker exited the oral cavity and always retracted more into the oral cavity. During drinking, however, the anterior tongue marker always leaves the oral cavity but only occasionally retracts fully into it (Figure 2). This is contrary to lapping in mammals with incomplete cheeks, in which the tongue always retracts into the oral cavity in successive cycles (Thexton et al., 1998). Whether protraction-retraction movements of the tongue are produced by movements of the tongue base or intrinsic regional deformations, or both, requires further investigation.

When observed without fluoroscopy, pigs appear to use suction to consume liquids, utilizing low amplitude, rhythmic jaw movements (Herring and Scapino 1973; Thexton et al., 1998; pers. obs.). However, according to Thexton et al., (1998), pigs utilize a combination of suction and lapping to transport the liquid bolus. The suction component of drinking may be created by the small amounts of jaw opening (i.e., decreasing Rz) that increase the volume of the oral cavity and creating a negative pressure within the oral cavity that draws water in. During drinking, we found that the anterior tongue does not undergo significant protraction-retraction, and the timing of its movement is highly variable. This suggests that the anterior tongue plays a minimal role in liquid ingestion. This is in contrast to the pronounced tongue protraction-retraction that occurs during lapping (e.g., Crompton and Musinsky 2011; Gart et al., 2015). Intrinsic tongue deformations may also contribute to the mechanics of sucking, particularly if shape changes occur in the intraoral region of the tongue. Compared to the significant and rapid oral cavity expansion that occurs during suction feeding for prey capture in aquatic vertebrates, as observed in many fish (e.g., Camp and Brainerd 2015; Lauder 1980a,b), the kinematics observed here suggest that pigs require only a small decrease in intraoral pressure for liquid to be drawn into the oral cavity. This is consistent with what has been proposed for the suckling mechanics of infant pigs (Thexton et al., 2004).

### Tongue-Jaw Coordination

We hypothesized that the timing of tongue protraction-retraction would be similar between behaviors primarily to avoid injury to the tongue. This is observed for the timing of maximum and minimum Tx of the posterior tongue marker, albeit with relatively high variability for drinking and for maximum retraction of the anterior tongue marker (i.e., minimum Tx value) during jaw closing. However, the timing of maximum Tx of the anterior tongue marker is significantly different between chewing and drinking (i.e., non-overlapping HPDs; Table 3). During chewing, maximum protraction occurs near the transition between SO and FO (75% of total gape cycle duration) with low variance for the anterior tongue marker. During this time, the tongue is collecting and repositioning the food along the tooth row. In contrast, during drinking, maximum protraction of the anterior tongue occurs at maximum gape (100% of total cycle duration, at the O2-closing transition), albeit with higher variance. This corresponds to the timing of tongue protraction observed during lapping in the cat (Thexton and McGarrick 1988). Thus, while there is some variability in protraction timing, the overall pattern is one that is consistent with retracting the tongue as the jaw closes, when there are both functional requirements associated with food or liquid transport as well as protection of the tongue as the teeth approximate. This also likely reflects fundamental properties of the motor control or orofacial movements: coactivation of jaw-opening with tongue-protruding muscles and jaw-closing with tongue-retraction muscles, which is known to occur across a variety of oral behaviors including mastication (e.g., Liu et al., 1993; Naganuma et al., 2001), licking (Travers et al., 1997), and infant suckling (Thexton et al., 1998). The fact that the motoneurons serving the groups of muscles coordinating tongue protrusion with jaw opening as well as tongue retraction with jaw closing share premotor neurons further supports our expectation (Stanek et al., 2014). Nevertheless, the more detailed analysis of these movements here demonstrates anteroposterior variation in tongue protrusion relative to jaw opening, a time when damage is unlikely to occur.

### Central Control of Chewing and Drinking Behaviors

These differences between drinking and chewing may provide insight into the changes that occur as infants shift from suckling to chewing solid foods and sucking for liquid ingestion. Mammalian infant suckling consists of negative pressure created by suction and/or physical expression of the teat (e.g., Herring 1985a; Thexton et. al., 2004). During both suckling and sucking in pigs, jaw opening appears to be the primary manner in which suction is created. During both behaviors, the tongue is outside the oral cavity for most of the cycle and anteroposterior tongue movements are small suggesting they contribute little to the creation of suction. As the tongue does not always return to the oral cavity and as the oral opening is continually submerged into the liquid, the small amount of tongue retraction is unlikely to form a liquid column as in lapping. Furthermore, both suckling and drinking appear to be bilaterally symmetrical (Lakars and Herring 1980), as compared to chewing which is unilateral, and has clear differentiation of sidedness, both in the jaw and the tongue movements (e.g., Abd-El-Malek 1955; Hiiemae and Palmer 2003). Chewing cycles occur at a lower frequency (3.1 vs 3.6 Hz) than drinking, and the frequency of drinking falls within the range of what is observed in infant pig suckling (3.5-4.4 Hz) (German et al., 1997). Therefore, drinking in adult pigs shares some common attributes with infant suckling.

Further investigation into the suckling CPG through the process of weaning would address how these movements are rhythmically controlled and modulated in relation to the development of the masticatory CPG. There is evidence for up to 6 CPGs present during early ontogeny (Barlow 2009; Nakamura et al., 2004; Tanaka et al., 1999), but how these relate to maturation or shifts in connections between different groups of premotor neurons and/or motoneurons controlling tongue and jaw movements throughout ontogeny is not understood. It appears that there is a shift from a cortical suckling area to a cortical masticatory area across ontogeny in the guinea pig (Iriki et al., 1988), reflecting developmental differences in sensorimotor centers associated with central pattern generation. Suckling rat pups show a motor pattern of nipple attachment that is very similar to that used for chewing whereas the motor pattern for rhythmic suckling from a nipple differs from the chewing motor pattern (Westneat and Hall 1992). In general, our results suggest that there are connections but also fundamental differences in the central control of sucking and chewing behaviors in pigs.

## CONCLUSIONS

The 3D kinematics of the jaw and tongue for chewing and drinking in pigs further our understanding of how these movements facilitate different oral behaviors. Drinking cycles were confirmed to be non-sided and instead only utilize two DoF: jaw pitch and anteroposterior translation. Chewing and drinking cycles were observed to have similar relative contributions of opening and closing to a standardized gape cycle, although with differing variability for each phase. Differences in tongue protraction-retraction magnitudes were observed, with larger magnitudes of movements observed during chewing. The timing of these movements indicates that some aspects of the tongue-jaw coordination pattern are different between these behaviors. Further, sucking in adults resembles infant suckling, including jaw opening to create suction and the anterior tongue positioned outside the oral cavity. Therefore, drinking cycles show characteristics of both chewing and infant suckling cycles, suggesting further research into the central control of different oral behaviors would provide valuable insight into the development of CPGs across different oral behaviors through ontogeny.

## ACKNOWLEDGMENTS

The authors would like to thank the Ohio University Laboratory Animal Resources staff for their help with animal husbandry. Dr. Andrew Niehaus at The Ohio State University College of Veterinary Medicine and Brooke Keener at the Holzer Clinic assisted with CT scanning of animals. We would also like to thank Volume Graphics GmbH for access to VGSTUDIOMAX 3.3 and Marissa O’Callaghan for her assistance with data collection and processing.

## DATA AVAILABILITY

All data used for this study, including metadata, CT scan data and the original unprocessed x-ray movies, are uploaded to the X-ray Motion Analysis Portal (http://xmaportal.org/webportal/).

## References

Abd-El-Malek, S. (1955). The part played by the tongue in mastication and deglutition. J. Anat. 89, 250–254.

Barlow, S. M. (2009). Central pattern generation involved in oral and respiratory control for feeding in the term infant. Curr. Opin. Otolaryngol. Head Neck Surg. 17 (3), 187–193.

Boughter, J.D., Mulligan, M. K., St John, S. J., Tokita, K., Lu, L., Heck, D. H., and Williams, R.W. (2012). Genetic control of a central pattern generator: rhythmic oromotor movement in mice Is controlled by a major locus near Atp1a2. PLoS One 7 (5), e38169.

Brainerd, E. L., Baier, D. B., Gatesy, S. M., Hedrick, T. L., Metzger, K. A., Gilbert, S. L., and Crisco, J. J. (2010). X-Ray Reconstruction of Moving Morphology (XROMM): precision, accuracy and applications in comparative biomechanics research. J. Exp. Zool. A Ecol. Genet. Physiol. 313A (5), 262–279.

Camp, A. L., and Brainerd, E. L. (2015). Reevaluating musculoskeletal linkages in suction-feeding fishes with X-Ray Reconstruction of Moving Morphology (XROMM). Integr. Comp. Biol. 55, 36–47.

Chen, X. (1998). The instantaneous center of rotation during human jaw opening and its significance in interpreting the functional meaning of condylar translation. Am. J. Phys. Anthropol. 106 (1), 35–46.

Crompton, A. W. (1989). The evolution of mammalian mastication. In Complex Organismal Functions: Integration and Evolution in Vertebrates (ed. D. B. Wake and G. Roth), pp. 23–40. John Wiley and Sons, Ltd.

Crompton, A. W., and Musinsky, C. (2011). How dogs lap: ingestion and intraoral transport in *Canis familiaris*. Biol. Lett. 7 (6), 882–884.

Cremers, J., and Klugkist, I. (2018). One direction? A tutorial for circular data analysis using R with examples in cognitive psychology. Front. Psychol. 9.

Davis, J. S. (2014). Functional morphology of mastication in musteloid carnivorans. PhD thesis, Ohio University, Athens, OH.

Dellow, P. G., and Lund, J. P. (1971). Evidence for central timing of rhythmical mastication. J. Physiol. 215 (1), 1–13.

Dotsch, C., and Dantuma, R. (1989). Electromyography and masticatory behavior in shrews (Insectivora). Prog. Zool. 35, 146–147.

Gart, S., Socha, J. J., Vlachos, P. P., and Jung, S. (2015). Dogs lap using acceleration-driven open pumping. Proc. Natl. Acad. Sci. 112 (52), 15798–15802.

German, R. Z., and Crompton, A.W. (1996). Ontogeny of suckling mechanisms in opossums (*Didelphis Virginiana)*. Brain Behav. Evol. 48 (3), 157–164.

German, R. Z., and Crompton, A. W. (2000). The ontogeny of feeding in mammals. In Feeding: Form, Function and Evolution in Tetrapod Vertebrates (ed. K. Schwenk), pp. 449–57. New York: Academic Press.

German, R. Z., Crompton, A. W., Levitch, L. C., and Thexton, A. J. (1992). The mechanism of suckling in two species of infant mammal: miniature pigs and longtailed macaques. J. Exp. Zool. 261 (3), 322–330.

German, R. Z., Crompton, A. W., Hertweck, D. W., and Thexton, A. J. (1997). Determinants of rhythm and rate in suckling. J. Exp. Zool. 278 (1), 1–8.

German, R. Z., Crompton, A. W., and Thexton, A. J. (2006). Ontogeny of feeding in mammals. In Feeding in Domestic Vertebrates: From Structure to Behavior (ed. V. Bels), pp. 50–61. Cambridge, NA: CABI.

Grossnickle, D. M. (2017). The evolutionary origin of jaw yaw in mammals. Sci. Rep. 7, 45094.

Herring, S. W. (1976). The dynamics of mastication in pigs. Arch. Oral Biol. 21, 473–480.

Herring, S. W. (1985a). The ontogeny of mammalian mastication. Am. Zool. 25, 339–349.

Herring, S. W. (1985b). Postnatal development of masticatory muscle function. Fortschritte Der Zoologie 30, 213–215.

Herring, S. W., and Scapino, R. P. (1973). Physiology of feeding in miniature pigs. J. Morphol. 141 (4), 427–460.

Herring, S. W., and Wineski, L. E. (1986). Development of the masseter muscle and oral behavior in the pig. J. Exp. Zool. 237, 191–207.

Herring, S. W., Raffertu, K. L., Liu, Z. J., and Marshall, C. D. (2001). Jaw muscles and the skull in mammals: the biomechanics of mastication. Comp. Biochem. Phys. A. 131, 207–219.

Hiiemae, K. M., and Palmer, J. B. (2003). Tongue movements in feeding and speech. Crit. Rev. Oral Biol. Med. 14 (6), 413–429.

Hiiemae, K. M., Thexton, A., and Crompton, A. W. (1978). Intra-oral food transport: the fundamental mechanism in feeding. In Muscle Adaptation in the Craniofacial Region. Monograph No. 8. Craniofacial Growth Series (ed. D. S. Carlson and J. McNamara), pp. 181–208. Ann Arbor: Ann Arbor Press.

Hylander, W. L. (1992). Functional anatomy. In The Temporomandibular Joint: A Biological Basis for Clinical Practice (ed. B.G. Sarnat and D.M. Laskin), pp. 60–92. Philadelphia: W.B. Saunders Co.

Iriarte-Diaz, J., Reed, D. A., and Ross, C. F. (2011). Sources of variance in temporal and spatial aspects of jaw kinematics in two species of primates feeding on foods of different properties. Integr. Comp. Biol. 51 (2), 307–319.

Iriki, A., Shuichi, N., and Nakamura, Y. (1988). Feeding behavior in mammals: corticobulbar projection is reorganized during conversion from sucking to chewing. Dev. Brain Res. 44, 189–196.

Knorlein, B. J., Baier, D. B., Gatesy, S. M., Laurence-Chasen, J. D., and Brainerd, E. L. (2016). Validation of XMALab Software for marker-based XROMM. J. Exp. Biol. 219 (23), 3701–3711.

Lauder, G. V. (1980a). Hydrodynamics of prey capture by teleost fishes. Biofluid. Mech. 2, 161–181.

Lauder, G. V. (1980b). The suction feeding mechanism in sunfishes (*Lepomis* spp.): an experimental analysis. J. Exp. Biol. 88, 49–72.

Lakars, T. C., and Herring, S. W. (1980). Ontogeny of oral function in hamsters (Mesocricetus Auratus). J. Morphol. 165 (3), 237–254.

Liu, Z. J., Masuda, Y., Inoue, T., Fuchihata, H., Sumida, A., Takada, K., and Morimoto, T. (1993). Coordination of cortically induced rhythmic jaw and tongue movements in the rabbit. J. Neurophysiol. 69 (2), 569–584.

Liu, Z. J., Shcherbatyy, V., Kayalioglu, M., and Seifi, A. (2009). Internal kinematics of the tongue in relation to muscle activity and jaw movement in the pig. J. Oral Rehabil. 36 (9), 660–674.

Lund, J. P., and Kolta, A. (2005). Adaption of the central masticatory pattern to the biomechanical properties of food. Int. Cong. Ser. 1284, 11–20.

Lund, J. P., and Kolta, A. (2006). Generation of the central masticatory pattern and its modification by sensory feedback. Dysphagia 21, 167–174.

Menegaz, R. A., Baier, D. B., Metzger, K. A., Herring, S. W., and Brainerd, E. L. (2015). XROMM analysis of tooth occlusion and temporomandibular joint kinematics during feeding in juvenile miniature pigs. J. Exp. Biol. 218 (16), 2573–2584.

Montuelle, S. J., Olson, R. A., Curtis H., Sidote, J. V., and Williams, S. H. (2018). Flexibility of feeding movements in pigs: effects of changes in food toughness and stiffness on the timing of jaw movements. J. Exp. Biol. 221 (2), jeb168088.

Montuelle, S. J., Olson, R. A., Curtis, H., Sidote, J. V., and Williams, S. H. (2019). The effect of unilateral lingual nerve injury on the kinematics of mastication in pigs. Arch. Oral Biol. 98, 226–237.

Montuelle, S. J., Olson, R. A., Curtis, H., Beery, S., and Williams, S. H. (2020a). Effects of food properties on chewing in pigs: flexibility and stereotypy of jaw movements in a mammalian omnivore. PLoS One 15 (2), e0228619.

Montuelle, S. J., Olson, R. A., Curtis, H., and Williams, S. H. (2020b). Unilateral lingual nerve transection alters jaw-tongue coordination during mastication in pigs. J. Appl. Physiol. 128 (4), 941–951.

Naganuma, K., Inoue, M., Yamamura, K., Hanada, K., and Yamada, Y. (2001). Tongue and jaw muscle activities during chewing and swallowing in freely behaving rabbits. Brain Res. 915 (2), 185–194.

Nakamura, Y., Katakura, N., and Nakajima, M. (1999). Generation of rhythmical ingestive activities of the trigeminal, facial, and hypoglossal motoneurons in in vitro CNS preparations isolated from rats and mice. J. Med. Dent. Sci. 46 (2), 63–73.

Nakamura, Y., Katakura, N., Nakajima, M., and Liu, J. (2004). Rhythm generation for food-ingestive movements. Prog. Brain Res. 143, 97–103.

Nozaki, S., Iriki, A., and Nakamura, Y. (1986). Localization of central rhythm generator involved in cortically induced rhythmical masticatory jaw-opening movement in the guinea pig. J. Neurophysiol. 55, 806–825.

Reis, P. M., Jung, S., Aristoff, J. M., and Stocker, R. (2010). How cats lap: water uptake by *Felis catus*. Science 330, 1231–1234.

Ross, C. F., Eckhardt, A., Herrel, A., Hylander, W. L., Metzger, K. A., Schaerlaeken, V., Washington, R. L., and Williams, S. H. (2007a). Modulation of intra-oral processing in mammals and lepidosaurs. Integr. Comp. Biol. 47 (1), 118–136.

Ross, C. F., Dharia, R., Herring, S. W., Hylander, W. L., Liu, Z. J., Rafferty, K. L., Ravosa, M. J., and Williams, S. H. (2007b). Modulation of mandibular loading and bite force in mammals during mastication. J. Exp. Biol. 210 (6), 1046–1063.

Ross, C. F., Baden, A. L., Georgi, J., Herrel, A., Metzger, K. A., Reed, D. A., Schaerlaeken, V., and Wolff, M. S. (2010). Chewing variation in lepidosaurs and primates. J. Exp. Biol. 213 (4), 572–584.

Ross, C. F., Iriarte-Diaz, J., Platts, E., Walsh, T., Heins, L., Gerstner, G. E., and Taylor, A. B. (2017). Scaling of rotational inertia of primate mandibles. J. Hum. Evol. 106, 119–132.

Smith, R. J. (1985). Functions of condylar translation in human mandibular movement. Am. J. Orthod. 88 (3), 191–202.

Stanek, E., Cheng, S., Takatoh, J., Han, B. X., and Wang, F. (2014). Monosynaptic premotor circuit tracing reveals neural substrates for oro-motor coordination. Elife 3, e02511.

Takahashi, T., Miyamoto, T., Terao, A., and Yokoyama, A. (2007). Cerebral activation related to the control of mastication during changes in food hardness. Neuroscience 145, 791–794.

Tanaka, S., Kogo, M., Chandler, S. H., and Matsuya, T. (1999). Localization of oral-motor rhythmogenic circuits in the isolated rat brainstem preparation. Brain Res. 821, 190–199.

Thexton, A. J., and McGarrick, J. D. (1988). Tongue movement of the cat during lapping. Arch. Oral Biol. 33 (5), 331–339.

Thexton, A. J., and Crompton, A. W. (1989). Effect of sensory input from the tongue on jaw movement in normal feeding in the opossum. J. Exp. Zool. 250 (3), 233–243.

Thexton, A. J., and McGarrick, J. D. (1989). Tongue movement in the cat during the intake of solid food. Arch. Oral Biol. 34 (4), 239–248.

Thexton, A. J., Crompton, A. W., and German, R. Z. (1998). Transition from suckling to drinking at weaning: a kinematic and electromyographic study in miniature pigs. J. Exp. Zool. 280 (5), 327–343.

Thexton, A. J., Crompton, A. W., Owerkowicz, T., and German, R. Z. (2004). Correlation between intraoral pressures and tongue movements in the suckling pig. Arch. Oral Biol. 49 (7), 567–575.

Travers, J. B., Dinardo, L. A., and Karimnamazi, H. (1997). Motor and premotor mechanisms of licking. Neurosci. Biobehav. Rev. 21 (5), 631–647.

Trulsson, M. (2007). Force encoding by human periodontal mechanoreceptors during mastication. Arch. Oral Biol. 52, 357–360.

Trulsson, M., and Johansson, R. S. (2002). Orofacial mechanoreceptors in humans: encoding characteristics and responses during natural orofacial behaviors. Behav. Brain Res. 135, 27–33.

Weijnen, J. (1998). Licking behavior in the rat: measurement and situational control of licking frequency. Neurosci. Biobehav. Rev. 22, 751–760.

Weijs, W. A., and De Jongh, H. J. (1977). Strain in mandibular alveolar bone during mastication in the rabbit. Arch. Oral Biol. 22 (12), 667–675.

Westneat, M. W., and Hall, W. G. (1992). Ontogeny of feeding motor patterns in infant rats: an electromyographic analysis of suckling and chewing. Behav. Neurosci. 106 (3), 539–554.

Withers, P. C., Cooper, C. E., Maloney, S. K., Bozinovic, F., and Cruz-Neto, A. P. (2016). Ecological and Environmental Physiology of Mammals. Oxford University Press.

