## Supplemental Tables for "Jaw Kinematics and Tongue Protraction-Retraction during Chewing and Drinking in the Pig"

**Supplemental Table 1**. Precision thresholds by individual and degree of freedom

| Individual ID | Tx (mm) | Ty (mm) | Tz (mm) | Rx (deg) | Ry (deg) | Rz (deg) |
|---|---|---|---|---|---|---|
| **Pig 20** | 0.474 | 0.919 | 0.505 | 0.548 | 0.199 | 0.380 |
| **Pig 21** | 0.0834 | 0.145 | 0.0707 | 0.149 | 0.170 | 0.416 |

**Supplemental Table 2**. Timing and variance of tongue Tx

| | Maximum Tx Timing (variance) | | Minimum Tx Timing (variance) | |
|---|---|---|---|---|
| | **Chew Cycles** | **Drink Cycles** | **Chew Cycles** | **Drink Cycles** |
| **Anterior** | 71.3% (2.4) | 1.2% (17) | 10.4% (12.7) | 25.8% (16.1) |
| **Posterior** | 78.8% (4.6) | 87.1% (18.5) | 43.2% (1.5) | 30.3% (10.0) |

Values are reported as a percentage of standardized cycle time.
