## Supplemental Figures for "Jaw Kinematics and Tongue Protraction-Retraction during Chewing and Drinking in the Pig"

**Supplemental Figure 1.** Schematic demonstrating interpretation of 95% highest posterior density intervals (HPDs). (A) An HPD of (25.0, 75.0), shown by the shaded area, indicates there is a 95% probability that the posterior mean is contained within this range. (B) An HPD of (75.0, 25.0) includes the cycle start, as indicated by the shaded region, as the start HPD position is higher than the end HPD position. Non-overlapping HPDs indicate a difference between behaviors whereas overlapping HPDs indicate a null hypothesis cannot be rejected. (A) and (C) have overlapping HPDs (from 25-35% of cycle duration), and therefore are not statistically different. The same is true for (B) and (C) as they also have overlapping HPDs (from 15-25% of cycle duration). (B) and (D) also overlap. Because the HPD for (D) is contained within (B), they are not statistically different. (A) and (D), and (C) and (D) do not have overlapping HPDs, and therefore have significantly different means.

**Supplemental Figure 2.** Graphs of the 6 DoF during 1.6 seconds of feeding (left) and drinking (right) from Individual 21. Translations are shown on the top and rotations are on the bottom. Shaded area represents the precision threshold about the sequence mean. * next to each shaded area indicates the DoF exceeds the precision thresholds.

**Supplemental Figure 3.** Both behaviors have a similar proportion of opening and closing phases. However, FO contributes more to total cycle duration for chewing cycles than O2 does for drinking cycles. Mean phase duration as a percentage of standardized gape cycle are shown for chewing (left) and drinking (right). Each wedge represents the proportion of mean total gape cycle duration for each phase. Chewing phases are: FC, fast close; SC, slow close; SO, slow open; and FO, fast open. Drinking phases are: C, close; O1, open 1; and O2, open 2.

**Supplemental Figure 4.** The average velocity of jaw pitch differs between chewing and drinking. Each datapoint represents the cycle duration and the corresponding maximum pitch (i.e., minimum Rz value) for a cycle. The least squares linear regression line and corresponding $R^2$ are shown for chewing cycles (open markers) and drinking cycles (solid markers). Individual 20 is represented by circles and Individual 21 by squares.

#### Supplemental Figure 1

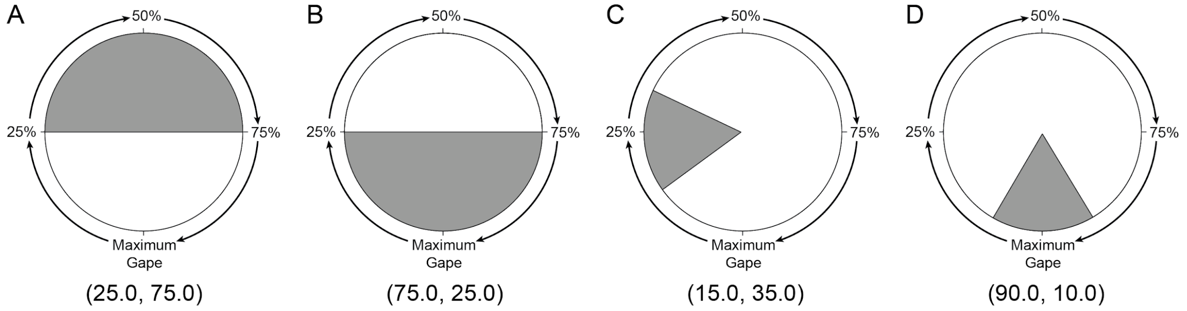

### Supplemental Figure 2

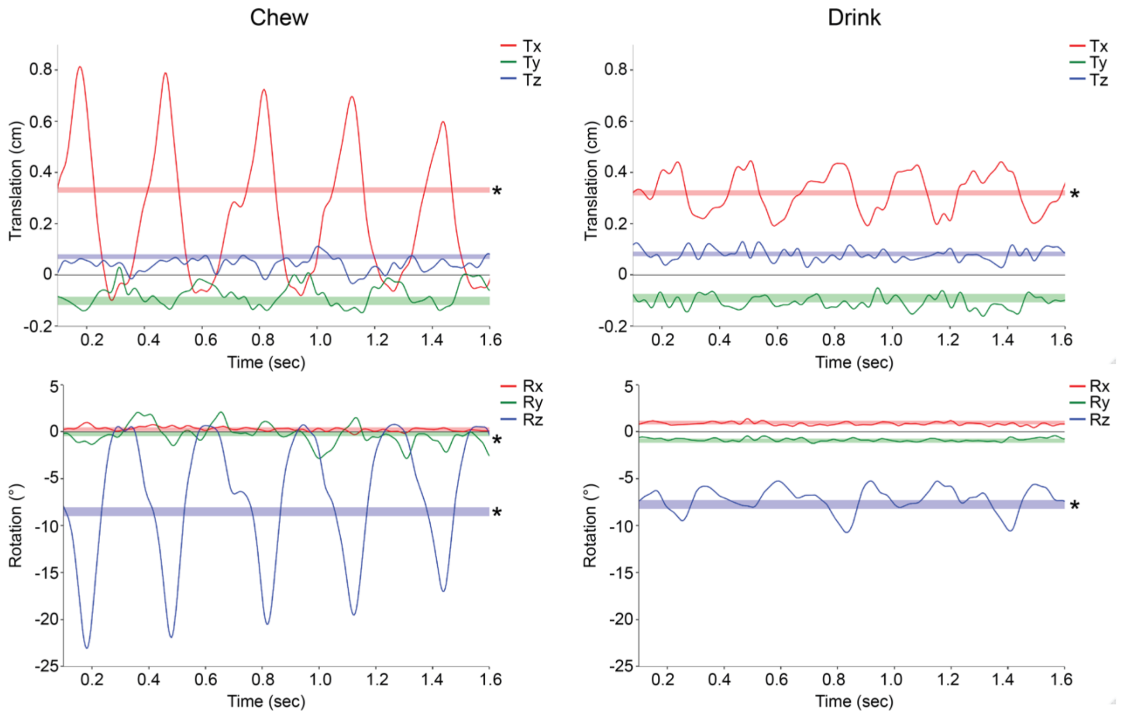

**Supplemental Figure 3**

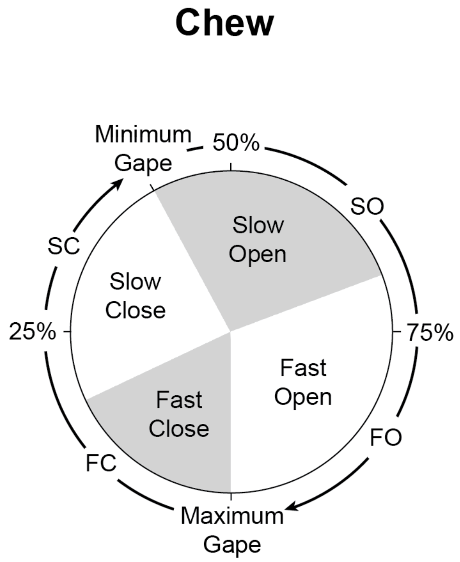

**Supplemental Figure 4**

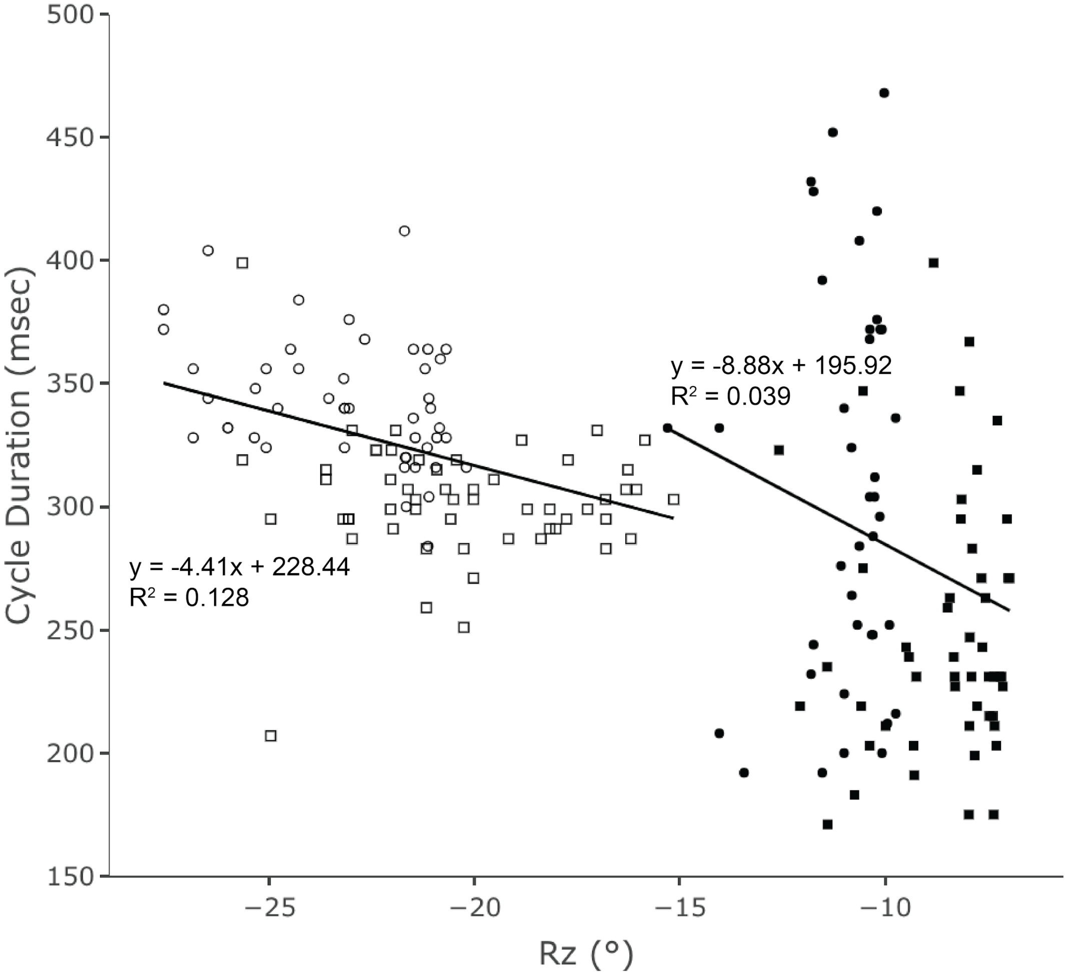
