## Supplemental Video Legends for "Jaw Kinematics and Tongue Protraction-Retraction during Chewing and Drinking in the Pig"

**Supplemental Video 1.** Fluoroscopy video of Individual 20 chewing a piece of apple.

**Supplemental Video 2.** Fluoroscopy video of Individual 20 drinking.

**Supplemental Video 3.** XROMM animation of Individual 20 chewing a piece of apple.

**Supplemental Video 4.** XROMM animation of Individual 20 drinking.
